# Spatiotemporal dissection of the cell cycle with single-cell proteogenomics

**DOI:** 10.1101/543231

**Authors:** Diana Mahdessian, Anthony J. Cesnik, Christian Gnann, Frida Danielsson, Lovisa Stenström, Muhammad Arif, Cheng Zhang, Rutger Shutten, Anna Bäckström, Peter Thul, Nathan H. Cho, Oana Carja, Mathias Uhlén, Adil Mardinoglu, Charlotte Stadler, Cecilia Lindskog, Burcu Ayoglu, Manuel D. Leonetti, Fredrik Pontén, Devin Sullivan, Emma Lundberg

## Abstract

Cellular division is a fundamental source of cell-to-cell variability, and studies of transcript and protein abundances have revealed several hundred genes that are regulated by the cell cycle^1–8^. However, none of these studies provide single-cell resolution of protein expression, leaving an incomplete understanding of cell-to-cell heterogeneity and the roles of cycling transcripts and proteins. Here, we present the first comprehensive map of spatiotemporal heterogeneity of the human proteome by integrating proteomics at subcellular resolution, single-cell transcriptomics, and pseudotime measurements of individual cells within the cell cycle. We identify that 17% of the human proteome displays cell-to-cell variability, of which 26% is correlated to cell cycle progression, and we present the first evidence of cell cycle association for 235 proteins. Only 15% of proteomic cell cycle regulation is due to transcriptomic cycling, which points to other means of regulation such as post-translational modifications. For proteins regulated at the transcript level, we observe a 7.7 hour delay between peak expression of transcript and protein on average. This spatially resolved proteomic map of the cell cycle has been integrated into the Human Protein Atlas and serves as a valuable resource for accelerating molecular studies of the human cell cycle and cell proliferation.

## INTRODUCTION

Cellular processes are driven by the presence and activity of specific proteins that are precisely regulated in time and space. The cell division cycle is tightly controlled at specific checkpoints^9^ by transcriptional regulation^10^, protein post-translational modification (PTM) feed-forward and feedback loops^11^, and protein degradation^12,13^. Dysregulation of the cell cycle has devastating consequences, such as uncontrolled cell proliferation, genomic instability^14^, and cancer^15,16^.

Studies of transcript and protein abundance variation between phases of the human cell cycle have shown that 400-1,200 genes^1–3^ and 300-700 proteins^4–8^ are regulated by the cell cycle. These studies have commonly been performed in bulk, with synchronized cell populations^1,2,17–20^. Hence, they have suffered from the disruptive nature of cell synchronization, known to alter gene expression^21^, perturb cell morphology and metabolism^22^. Synchronization of cells also precludes the discovery of expression changes within cell cycle phases. Single-cell RNA sequencing has allowed the analysis of transcriptional changes without the need for synchronization and enables the discovery of more cell cycle regulated genes^23,24^. However, due to technological limitations, it has not been possible to systematically study single cell variations at the proteome-wide level.

Here, we report on a systematic characterization of temporal expression patterns of RNA and protein with single-cell resolution in unsynchronized human cells by leveraging the antibody resource^26^ and image collection of the Human Protein Atlas (HPA)^25^. This represents the first spatially resolved map of human proteome dynamics during the cell cycle obtained by imaging proteogenomics. We provide knowledge of whether these events are regulated by transcript cycling by performing single-cell RNA sequencing, and we analyze the contributions of protein stability and PTMs to cell cycle regulation by using bulk MS proteomic data.

We find that a large portion of the human proteome (18%, 2,292/12,390 proteins) displays cell-to-cell heterogeneity in terms of level of expression. We performed a targeted single-cell proteomic imaging and single-cell transcriptomic analysis (*i.e.*, imaging proteogenomics) of the 1,219 proteins expressed in FUCCI^27,28^ U-2 OS cells and found that 26% (321/1,219) of these proteins had variation correlating with cell cycle progression. Our systematic analysis identifies 235 novel cell cycle dependent (CCD) proteins and shows that only a small portion (15%) of proteomic cell cycle regulation can be attributed to transcript cycling. Other properties differentiate the transcriptionally and non-transcriptionally regulated protein groups. We see tighter PTM regulation for cycling proteins and looser PTM regulation for proteins with cycling transcripts. In contrast, transcriptionally regulated CCD proteins are significantly less stable, indicating a larger role of unfolding and degradation in their cell cycle regulation. Finally, we show that cycling proteins are generally enriched in proliferating tissues, and several may have oncogenic or anti-oncogenic functions.

## RESULTS

### Proteomic heterogeneity of single cells

The HPA Cell Atlas aims to localize human proteins at a subcellular level using immunofluorescence and confocal microscopy^25^. This high-resolution image collection visualizes protein expression in a variety of cell lines, always unsynchronized and in log-phase growth, and provides an unprecedented resource to explore protein expression heterogeneity at single-cell level. Out of the 12,390 proteins mapped in the HPA Cell Atlas, 2,292 (19%, Supplementary Table 1) show cell-to-cell heterogeneity in terms of expression level or spatial distribution (Methods). CCNB1 is a well known cell cycle regulator^29^ that exemplifies such variation in abundance, and HMGSC1 displays variation in abundance and spatial localization between the nucleus and the cytosol (Fig. 1A). Out of these 2,292 proteins, 66% showed similar cell-to-cell variations in more than one human cell line (Supplementary Table 1), as exemplified by RACGAP1 (Fig. 1B). This suggests that these proteome variations might be controlled by conserved regulatory mechanisms. In this study, we investigate to what extent these observed protein variations represent temporally controlled expression patterns correlated to cell cycle progression.

**Figure 1:**
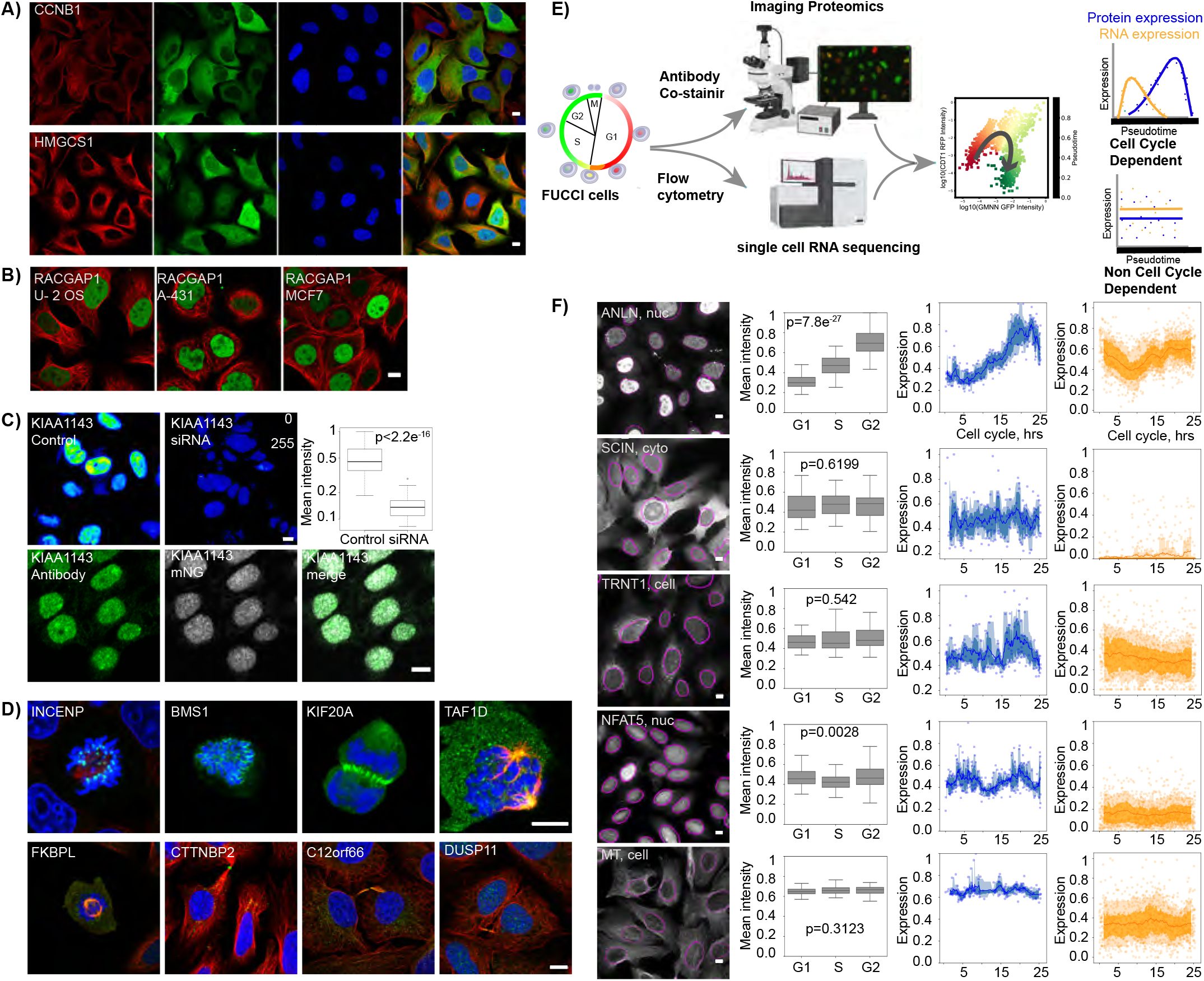
Temporal dissection of cell-to-cell heterogeneity using imaging proteogenomics. In A-D, the target protein is shown in green, microtubules in red, and the nucleus in blue. Scale bars in A-F represent 10μm. A: Examples of immunofluorescent images of U-2 OS for proteins with observed cell-to-cell heterogeneity in protein abundance (CCNB1) and spatial distribution (MRTO4). B: The RACGAP1 protein shows consistent cell-to-cell heterogeneity in several cell types (U-2 OS, A-431 and MCF7). C: Examples of antibody validation. The specificity of the anti-KIAA1143 antibody is validated with siRNA-mediated gene silencing (top) that resulted in significantly lower staining intensity. The specificity is also validated by co-localization with mNG-tagged KIAA1143. D: Example images of proteins localized to mitotic substructures: INCENP to kinetochores, BMS1 to mitotic chromosomes, KIF20A to cleavage furrow, TAF1D and FKBPL to mitotic spindle, CTTNBP2 to midbody ring, and C12orf66 and DUSP11 to cytokinetic bridge. E: Schematic overview of the single-cell imaging proteogenomic workflow. U-2 OS FUCCI cells express two fluorescently-tagged cell cycle markers, CDT1 during G1 phase (red) and Geminin during S and G2 phases (green); these markers are co-expressed during the G1-S transition (yellow). By fitting a polar model to the red and green fluorescence intensities, a linear representation of cell cycle pseudotime is obtained. Independent measurements of RNA and protein expression are compared after pseudotime alignment of individual cells. F: Examples images of the proteins ANLN, CPSF6, and NFAT5 with alpha-tubulin (MT) as negative control. The boxplot shows a mock-up bulk proteomic experiment. Single cell pseudotime expression profiles are shown for protein (blue) and RNA (orange).

The antibodies used in this study are validated in the HPA standard pipeline (Supplementary Table 1 and 2). In addition, we tagged a large number of proteins at endogenous levels with mNeonGreen using CRISPR/Cas9, and validated antibodies by gene silencing using siRNA and through a paired antibody approach (Methods, Fig. 1C, Extended Data Fig. 1, Supplementary Table 3).

### Proteins in mitotic structures

A protein can be directly associated with the cell cycle if it localizes to a mitotic structure. We found 230 proteins mapped to one or several mitotic structures (92 to cytokinetic bridge, 65 to mitotic chromosome, 39 to mitotic spindle, 35 to midbody, 14 to midbody ring, 4 to kinetochores, 3 to cleavage furrow; Fig. 1D). Among these, 99 were not previously associated with the cell cycle (Methods; Supplementary Table 4), for example C12orf66 and DUSP11 in the cytokinetic bridge, and BMS1 on mitotic chromosomes. Novel proteins identified at the mitotic spindle include TAF1D involved in transcription, and FKBPL, which is known to be an anticancer protein that is crucial for response to radiation stress^30^.

### Interphase proteogenomics in single cells

To determine if observed cell-to-cell variations correlate with interphase progression, we performed targeted single-cell imaging proteogenomics analysis using FUCCI^27,28^ U-2 OS cells (Fig. 1E). Of the 2,292 proteins identified to show cell-to-cell variability, the 1,219 proteins expressed in U-2 OS cells were selected for in-depth analysis of protein localization and abundance using immunofluorescence; single-cell transcriptomic analysis was performed by single-cell RNA sequencing. These proteins were measured to have a mean fold-change of 6.89 between the highest and lowest expressing cells. We also determined that 992 proteins (81%) had unimodal and 227 proteins (19%) had bimodal population intensity distributions (Extended Data Fig. 2). The pseudotemporal position of each cell in interphase was measured using FUCCI markers (Fig. 1E, Extended Data Fig. 3). Examples of this analysis are given in Fig. 1F. ANLN, a well-characterized cell cycle regulator^31^ increased in nuclear abundance during cell cycle progression, as verified by time-lapse imaging, and showed a similar temporal profile of RNA expression; in contrast, HPSE, an enzyme that cleaves heparan sulfates, revealed revealed variation in the nucleus that did not correlate with the cell cycle (Extended Data Fig. 4). We aggregated single-cell measurements by cell cycle phase to simulate a bulk experiment, and 73 CCD proteins detected at the single cell level were not detectable in bulk phases, such as TRNT1. NFAT5 also provides an example where single cell resolution allows us to observe a second harmonic CCD oscillation that is not reflected in bulk. Staining of alpha-tubulin in all samples served as a negative control, with no significant variation of expression during cell cycle progression (Extended Data Fig. 5).

### Discovering cell cycle regulated proteins

Based on this analysis, we identified 321 out of 1,219 proteins (26%) to have variance in expression levels temporally correlated to cell cycle interphase progression (Methods, Fig. 2A, Supplementary Table 2, Extended Data Fig. 2). It is noteworthy that most proteins analyzed (71%) showed cell-to-cell variations that were largely unexplained by cell cycle progression (hereafter denoted ‘non-CCD’), which opens up intriguing avenues for further exploration of the stochasticity or deterministic factors that govern these variations. Another key finding is that the variance of many cell cycle regulated proteins is only partially explained by the cell cycle, particularly for the newly identified ones. We hypothesize that these proteins are deterministically controlled by other cellular mechanisms, which opens the door to follow up on the role of the cross-talk of various signaling pathways in cell cycle regulation.

**Figure 2:**
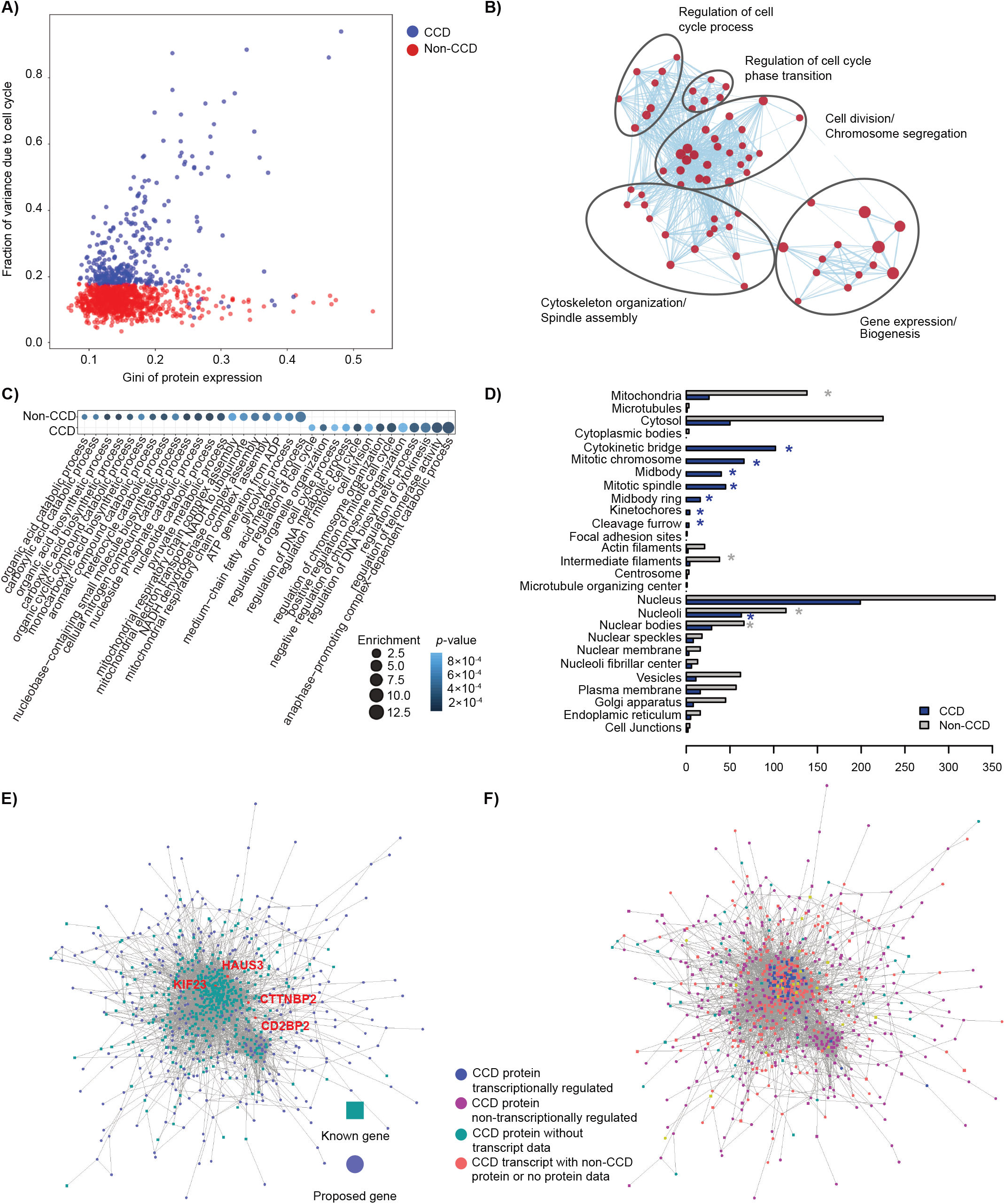
Properties of proteins with cell-to-cell variability. A: Scatter plot of percentage explained variance and Gini index for each investigated protein. Blue color indicates a significant cell cycle dependence (CCD), and red color indicates variability independent of the cell cycle (non-CCD). B: Gene ontology enrichment analysis for CCD proteins shows significantly enriched terms for the BP domain. Each circle represents a GO term, and line width corresponds to the number of genes that overlap between the two connected gene sets. Similar terms are grouped and labeled in the figure. C: Gene ontology enrichment analysis for CCD and non-CCD enzymes shows enrichment for BP domain terms. CCD enzymes are enriched for cell cycle related functions, whereas non-CCD enzymes are enriched for basic metabolic functions. D: Bar plot showing the subcellular localizations of CCD proteins (blue) and non-CCD proteins (grey). E: Protein-protein interaction (PPI) network of CCD proteins using the STRING^35^ database. The protein with a known association to the cell cycle (by GO term, Reactome pathway, or Cyclebase phenotype) are shown in green squares. F: PPI network of CCD transcripts and proteins using the STRING database.

Out of the 1,424 proteins, found in mitotic structures (230 proteins) or analyzed in interphase (1,219 proteins), 541 were CCD proteins (322 in interphase, 230 in mitotic structures, and 10 in both), and only 235 (43%) had a known association to the cell cycle by GO term^32^, Reactome pathway^33^, or Cyclebase phenotype^34^ (hereafter denoted ‘known CCD’. The remaining 306 proteins (57%), had no documented association to the cell cycle (hereafter denoted ‘novel CCD,’ Supplementary Tables 2 and 4). The set of genes corresponding to CCD proteins (known and novel) was highly enriched for GO terms related to cell division (Fig. 2B). Interestingly, the genes corresponding to non-CCD variable proteins were not enriched for any GO biological process (BP) terms. When limiting the GO enrichment analysis to enzymes, terms related to basic metabolic functions (e.g., ATP generation) or entire metabolic pathways (e.g., glycolysis) were overrepresented for the non-CCD variable proteins (Fig. 2C). This shows that the identified CCD proteins are indeed involved in cell cycle processes, whereas the proteins not correlated to the cell cycle are likely involved in a variety of different biological processes including metabolism.

The high resolution of our analysis allows us to study the role of subcellular localization in cell cycle regulation. We found significant differences in the localization of proteins that show CCD or non-CCD expression (Fig. 2D). Proteins with variations independent of the cell cycle were enriched in intermediate filaments, nucleoli, nuclear bodies, and mitochondria, whereas CCD proteins were enriched in nucleoli and mitotic structures, constituting 39% of the CCD proteins (*p* < 0.01 for both enrichments, binomial one-sided test, mapped proteome as background, significant enrichments marked as asterisks in Fig. 2D). Almost half (43%) of the CCD proteins resided in the nuclear compartment (3% nuclear speckles, 9% nuclear bodies, 22% nucleoli and 66% nucleus), not surprisingly given that one of the main functions performed in the nucleus is the replication of DNA during the cell cycle.

Besides identifying novel CCD proteins, this study also for the first time establishes exact temporal expression profiles for many proteins known to be important for proliferation.

### Expanded network of cell cycle proteins

To investigate whether the novel CCD proteins are connected to known cell cycle proteins, we analyzed their protein-protein interactions using the STRING database^35^ and found significantly more interactions within this dataset than for random datasets of similar sizes (Lambda calculations PPI enrichment p-value < 1×10^−16^; 14,539 interactions; 4,591 expected interactions). This indicates that the novel CCD proteins are likely involved in related biological processes as known CCD proteins. The known CCD proteins (most often regulated at the transcript level) were tightly clustered in the core of the network, whereas the novel CCD proteins formed an extended network (Fig. 2E-F). For instance, KIF23 is essential for microtubule bundling during cytokinesis and is known to oscillate temporally in the nucleus during the cell cycle^36^; we show that among KIF23 interactors are both known cell cycle regulators and novel CCD proteins. One such protein is CTTNBP2, which was localized to the midbody ring (Fig 1D) and further interacts with HAUS3 that is also known to be required for cytokinesis. These interactions imply that CTTNBP2 is also involved in cytokinesis. Another interesting set of interactions are those of CD2BP2 (Fig 2E, 5E), whose expression peaks in late G2, with a cluster of proteins needed for chromosome segregation located at mitotic chromosomes, implicating CD2BP2 in the transition or regulation of mitosis.

### RNA cycling underlies few cycling proteins

We investigated whether the regulation of cell cycle proteins could be attributed to transcript cycling by analyzing single cell RNA-sequencing data (Methods, Fig. 1E). The cycling of these cells can clearly be seen in the UMAP projection that forms a cycle reflecting interphase progression (Fig. 3A, Extended Data Fig. 6A-B). The RNA velocity streams overlaid onto this projection illustrate CCD post-transcriptional regulation, since the velocities are calculated using the ratios of spliced to unspliced RNA, and they indicate larger magnitude of changes in G1 and G2 and smaller changes during DNA replication in S-phase. By overlaying the measured pseudotime, we can also validate that the FUCCI model accurately reflects interphase progression for RNA. We identified 402 of 13,450 protein-coding genes analyzed (3%) to have variance in expression levels correlated to cell cycle progression (Extended Data Fig. 6C, Supplementary Table 5). This includes 86% of known CCD transcripts^37^. Interestingly, only 72 were also detected as CCD proteins (15%). Of the CCD proteins, 85% did not have CCD transcripts. This small overlap of CCD proteins and transcripts is corroborated by external RNA datasets^19,38^ and indicates that the temporal dynamics of proteome regulation may be largely maintained at a translational or post-translational level.

**Figure 3:**
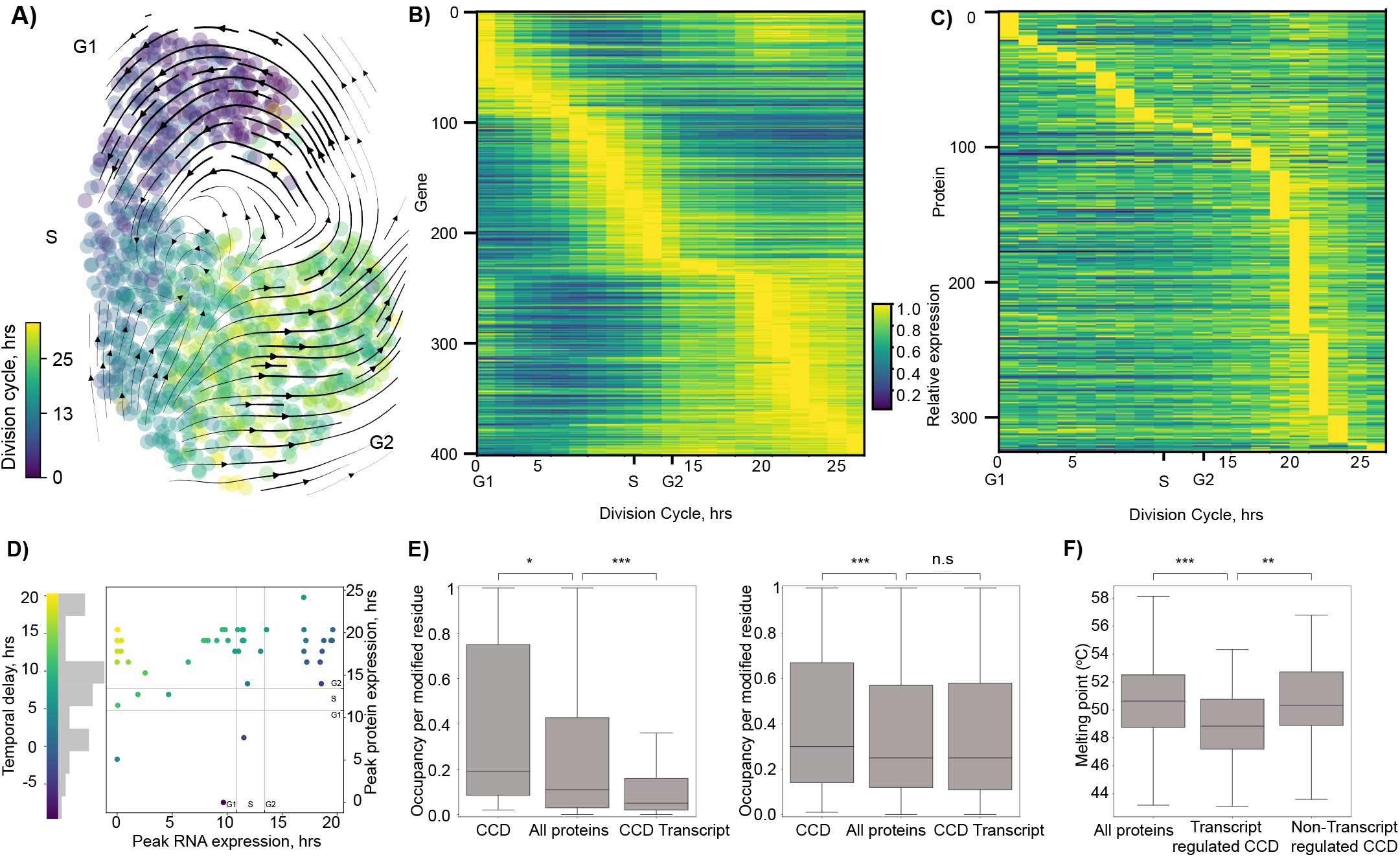
Proteogenomic dissection of cell cycle related variations. A: Dimensionality reduction of single-cell RNA sequencing data using UMAP shows a cycle correlated with pseudotime (color scale). RNA velocity moments are displayed as arrows that confirm the directionality of cycling, show faster movement through G1 (top) and G2 (bottom right), and slower progression through S phase (left). B: Heat map sorted by time of peak RNA expression for CCD transcripts, showing the relative RNA expression levels across the cell cycle. C: Heat map sorted by time of peak protein expression for CCD proteins, showing the relative protein expression levels across the cell cycle. D: The delay between peak RNA and protein expression is shown as a color scale and histogram. Each of the 72 genes with both CCD proteins and CCD transcripts are shown in a scatter plot, illustrating that most RNA peak in G1, and most proteins peak in G2 with a median delay of 7.7 hrs. E: The occupancy of PTMs for peptides measured in bulk (left) and phospho-enriched (right) MS proteomic experiments. Significantly higher occupancy is observed for CCD proteins than all proteins, indicating tighter regulation of a few dozen abundant PTMs including phosphorylations; significantly lower occupancy for PTMs on proteins with cycling transcripts in bulk but not in phospho-enriched data indicates looser regulation of a few dozen abundant PTMs excluding phosphorylations. F: Melting points of proteins from thermal profiling MS proteomic data indicate that transcript regulated CCD proteins melt at significantly lower temperatures, indicating lower stability that may aid protein degradation necessary for proteomic cycling in response to transcript cycling.

### Temporal delay between RNA and protein

To study the temporal dynamics of the cell cycle transcriptome and proteome, we sorted RNAs (Fig. 3B) and proteins (Fig. 3C) based on their time of peak expression. G1 is the longest period of the cell cycle, in which most RNAs are known to peak in expression (44%, 178/402, in this study; G1 10.8h; S-transition 2.6h; G2 11.9h in U-2 OS FUCCI cells), as corroborated by previous studies^3,39^. However, most proteins (235/321, 75%) peak towards the end of the cell cycle corresponding to late S and G2. We observed a 7.7 hr delay on average between peaks of RNA and protein expression for the 50 genes with both CCD proteins that are transcriptionally regulated (Fig. 3D, Supplementary Table 6). This is the first report of the delay between transcript and protein expression with single-cell resolution and is comparable to the temporal delay in circadian cycling^40^.

### PTMs and stability of cycling proteins

PTMs are important for cell cycle signaling, as exemplified by the classic cyclin dependent kinases (CDKs)^11^. We used bulk and phospho-enriched mass spectrometry (MS) proteomic data for U-2 OS cells to survey 63 types of abundant PTMs and deeply profile three types of phosphorylation on proteins regarding the cell cycle. The occupancy at each PTM site was calculated by comparing the number of modified and unmodified peptides observed at each site (Methods).

We observed higher occupancies for PTM sites on CCD proteins when compared to the average of all PTMs in both experiments, which points to tighter control of PTMs for these proteins across interphase. The 39 PTM sites observed on 23 CCD proteins in bulk had significantly higher occupancies when compared to the average of all PTMs (*p* = 0.018), as did the 1275 PTM sites on 124 CCD proteins in phospho-enrichment (*p* = 1.1×10^−8^; one-sided Kruskal-Wallis tests; Fig. 3E; Supplementary Tables 7-8). The strongly CCD protein DUSP19 (Fig. 4A) provides an example of tight PTM regulation in bulk analysis with a high occupancy citrullination at position 22 (28% of the 108 peptides crossing the position were modified).

**Figure 4:**
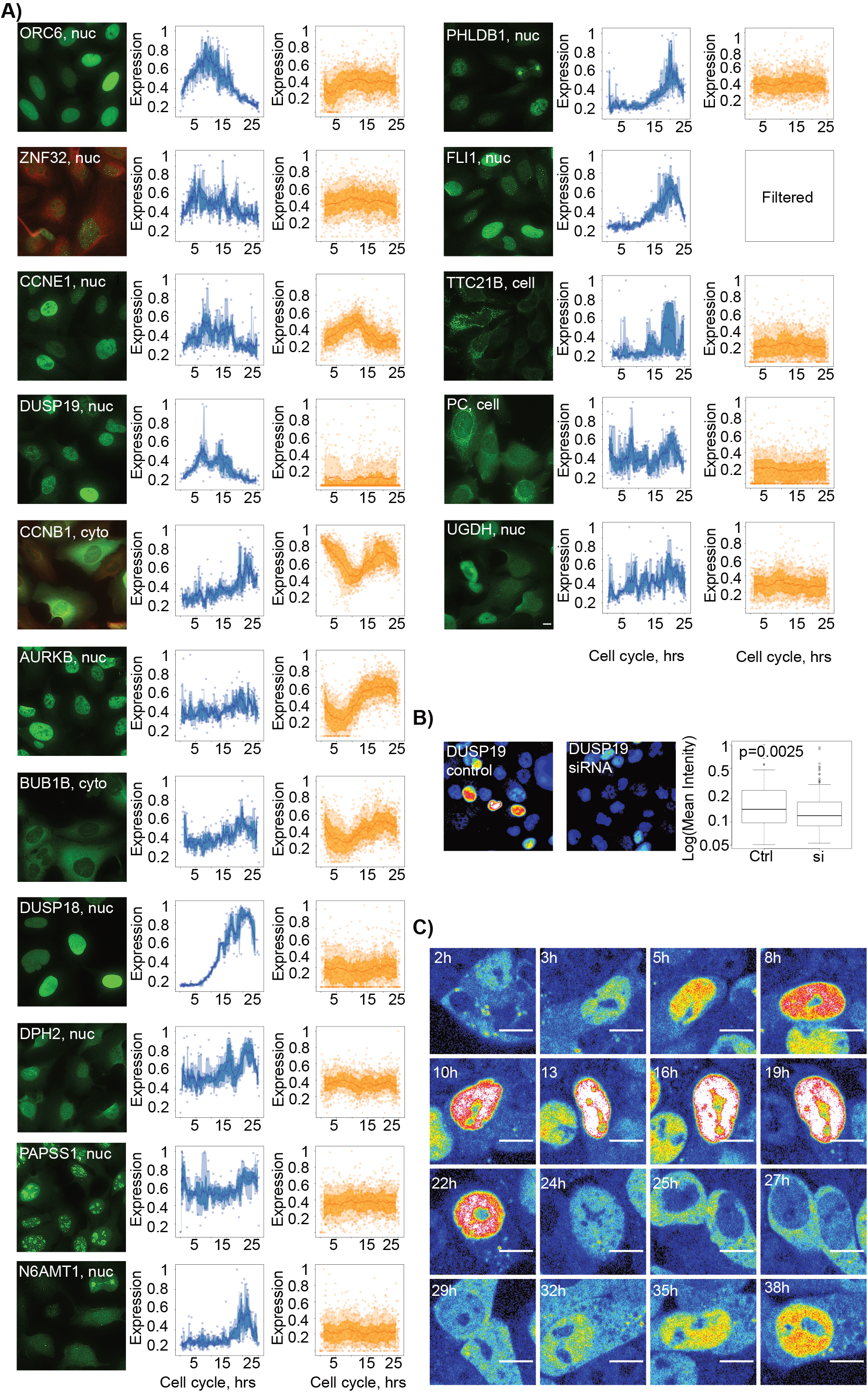
Cell cycle temporal expression profiles. A: Examples of temporal expression profiles for single-cell protein (blue) and RNA expression (orange). The compartment for which the protein abundance was measured is shown in brackets (nuc: nucleus, cyto: cytoplasm, cell: entire cell). The proteins of interest are shown in green, and microtubules are shown in red. B: Timelapse for mNG-tagged UGDH protein demonstrates cytoplasmic localization in G1 and translocation to the nucleus and an increase in expression during cell cycle progression. This live cell imaging data also corroborates the peak protein expression observed in G2 with immunofluorescence. C: The specificity of the antibody targeting DUSP19 is validated with siRNA-mediated gene silencing (si), resulting in a significant reduction of the staining intensity from a control (Ctrl).

**Figure 5:**
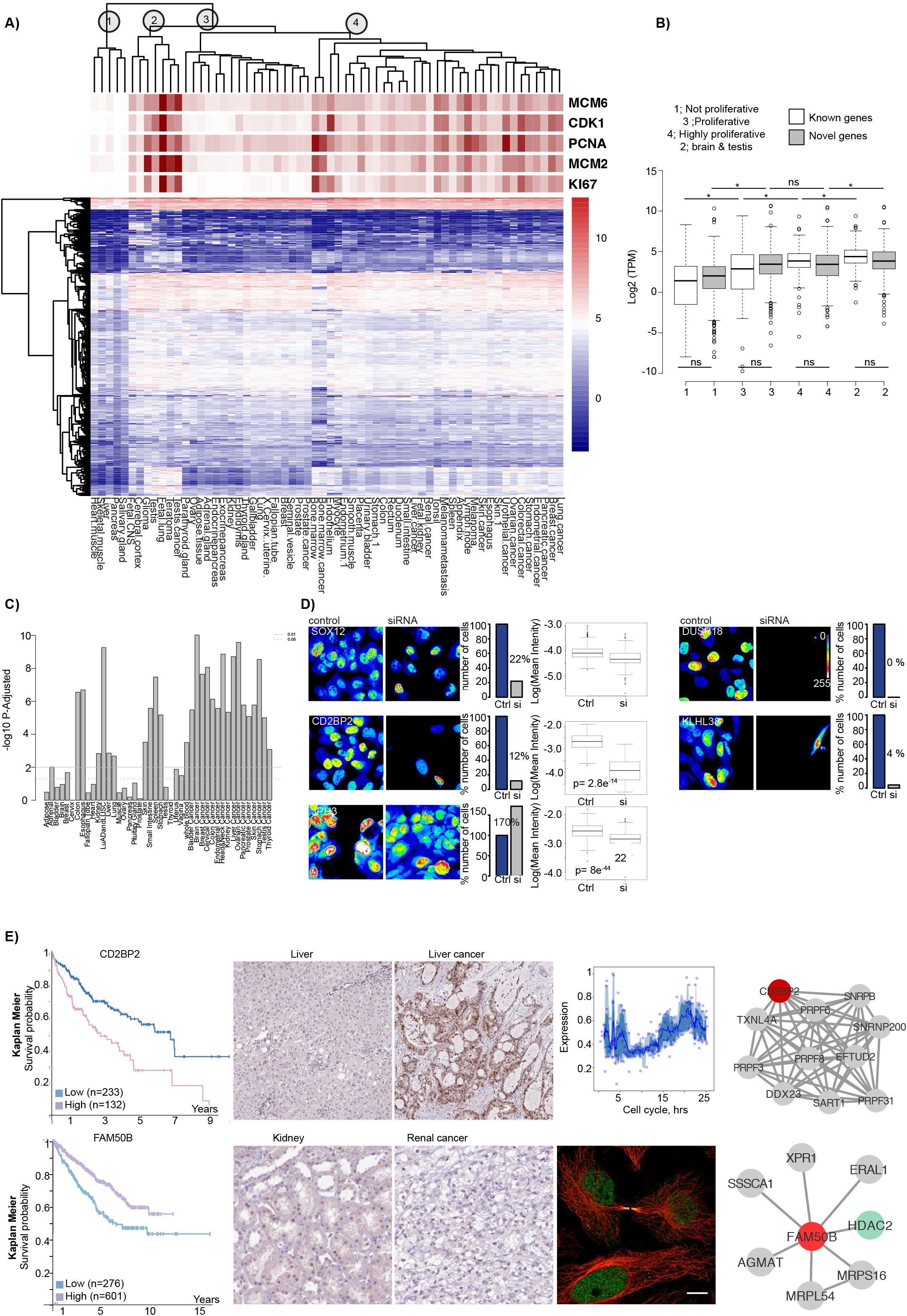
Functional associations for novel cell cycle regulated proteins. A: Hierarchical clustering of bulk transcript expression (log-transformed TPM values) for CCD proteins derived from RNA sequencing of various normal and cancer tissue types. The expression levels of the proliferation markers MCM6, CDK1, PCNA, MCM2 and KI67 are highlighted on top as a general measure of the proliferative activity of the tissues. Four clusters are identified: (1) contains normal tissues with low proliferative activity, (2) contains cerebral tissues with testis, (3) contains mostly normal tissues with midrange expression level of the proliferation markers and (4) contains tissues with high expression of the proliferation markers, including tumors. B: Box plots of the average transcript level for known and novel CCD proteins, respectively, for the four different clusters from A. C: Bar plot showing the significance of novel CCD proteins having shorter path distances to known CCD proteins than random in gene co-expression networks in various normal and cancer tissues. D: Silencing of CCD proteins SOX12, DUSP18, CD2BP2, KLHL38 and JPH3. Immunofluorescence images of the control and siRNA samples observed with LUT show variation in protein expression from low (blue) to high intensity (white). Bar plots show the differences in cell counts for control (Ctrl) and siRNA (si) samples, and boxplots show the significant decrease of the measured intensity (too few cells were observed in DUSP18 and KLH38 siRNA samples to make this comparison). E: Kaplan-Meier plots showing the correlation between survival and gene expression (FPKM) for FAM50B (top panel) and CD2BP2 (bottom). Higher expression of FAM50B was associated with longer survival (favorable) in renal cancer, and higher expression of CD2BP2 was associated with shorter survival (unfavorable) in liver cancer. Immunohistochemical images (target protein: brown, nuclei: blue) show lower expression of FAM50B in renal cancer than normal kidney and higher expression of CD2BP2 in liver cancer than normal liver. The protein-protein interaction network for FAM50B shows interaction with HDAC2, which is involved in the regulation of cell cycle progression, whereas CD2BP2 interacts with genes involved in splicing processes.

We also observed lower occupancies of PTMs on proteins with cycling transcripts than the average for all PTMs with the exception of phosphorylations, indicating looser regulation of these PTMs during interphase for proteins that are transcriptionally regulated. The 325 PTM sites on 60 proteins with cycling transcripts in bulk had significantly lower occupancies when compared to the average for all PTMs (*p* = 1.6×10e^−11^); this was not the case for the 2,657 PTM sites on 223 proteins with cycling transcripts in phospho-enrichment (*p* = 0.62; one-sided Kruskal-Wallis tests; Fig. 3E). The looser regulation of the 27 types of abundant PTMs excluding the three types of phosphorylation points to the larger role of transcriptional regulation for these proteins.

Protein degradation also plays a fundamental role in the cycling of proteins. Proteins are unraveled by the proteasome to initiate degradation, and so proteins that are more susceptible to unraveling should be more quickly degraded without stabilization. We investigated this phenomenon using protein melting points averaged from MS proteomic thermal profiling of A549, HEK293, and HepG2 cells^41^ (Supplementary Table 9). Lower melting points correspond to lower thermodynamic stability and lower thresholds for unraveling and degradation. We observed significantly lower stability for the 58 transcript-regulated CCD proteins than all 9,106 proteins and 237 non-transcriptionally regulated CCD proteins (*p* = 0.00030 and 0.0025, Fig. 3F). This may be important for degradation of these cycling proteins in response to transcript cycling. As the transcript disappears, proteins are no longer being actively produced, and the lower thermodynamic stabilities of these proteins may aid in allowing degradation rates to dominate, eliminating these proteins to complete the cycle of protein expression.

### Spatiotemporal partitioning of proteins

The temporal expression analysis enabled identification of proteins that share highly similar temporal patterns with well-known cell cycle regulators but have no prior association to the cell cycle (Fig. 3C,Fig. 4A). For instance, in the G1 group, well-known CCD proteins such as ORC6, required for the cell entry into S phase^42^, and MCM10, required for DNA replication^43^, were identified to have similar patterns as the novel CCD protein ZNF32, whose overexpression has been associated with a shorter survival time in lung adenocarcinoma cells^44^.The group peaking at the end of G1 includes known proteins such as CCNE1, along with the novel CCD protein DUSP19, a phosphatase whose depletion results in increased mitotic defects^45^. The specificity of the antibody targeting DUSP19 was validated with gene silencing (Fig. 4B). Time-lapse imaging can be used to verify the antibody based findings (Extended Data Fig. 7). The dehydrogenase UGDH was identified as a novel CCD protein that translocates from the cytosol to the nucleus in early G1, followed by a gradual increase in nuclear abundance until peak expression in G2, as observed with both immunofluorescence and live cell imaging of mNG tagged protein (Methods, Fig. 4C). UGDH produces precursors to glycosaminoglycans and proteoglycans of the extracellular matrix (ECM). Reduced UGDH expression inhibits glioblastoma growth and reduces ECM protein expression^46^. Future functional studies are needed to understand a potential non-canonical function for this enzyme in the nucleus and its contribution to the pro-proliferative function. The G2 group includes known proteins, such as CCNB1, AURKB (Extended Data Fig. 7A-B) and BUB1B. Novel CCD proteins in this group include the phosphatase DUSP18 (Fig. 1F); the transcription factor NFAT5 (Fig. 1F); an estrogen sulfating enzyme PAPSS1, for which overexpression has been reported to affect proliferation^47^; the methyltransferase N6AMT1; the uncharacterized protein PHLDB1; the enzyme DPH2; and transcription factor FLI1.

Shared subcellular localization can serve as another important association to known cell cycle proteins. For example, the mitochondrial protein pyruvate carboxylase (PC) was peaked in G2 (20.3 hrs) and shares the same subcellular location and temporal expression pattern as TTC21B (19.1 hrs). PC is involved in cell proliferation and gluconeogenesis and is upregulated in several types of cancer^48^, whereas TTC21B has no previously described association to cell proliferation. In this manner, we associated novel and known cycling proteins with similar temporal profiles in organelles such as the cytosol, nucleus, nucleoli and the Golgi apparatus.

Knowledge about the spatial and temporal expression profiles in relation to known cell cycle regulators is valuable for gathering a deeper causal understanding of the molecular effects of cell cycle progression and cellular proliferation.

### Associations with proliferative activity

To glean insight into whether the novel CCD proteins are functionally important for cellular proliferation, we analyzed bulk mRNA expression data across cohorts of normal and cancer tissue. Hierarchical clustering of the transcript data provided four major clusters: normal tissues with low proliferative activity, cerebral tissues together with testis as an outlier, normal tissues of moderate proliferative capacity, and cancers or normal tissues with high proliferative activity (Fig. 5A, Extended Data Fig. 8). Gene expression of the novel CCD proteins was significantly higher in proliferative tissues than non-proliferative tissues, indicating their importance for cellular proliferation (*p* = 2.9×10^−14^, one-sided Kruskal-Wallis test; Fig. 5B).

Using gene co-expression networks^49^, we show that novel CCD genes had significantly shorter paths to known cell cycle genes in all cancer tissues and normal proliferative tissues, whereas there was no significant decrease of path lengths in low-and non-proliferating tissues (Fig. 5C; one-sided Kolmogorov-Smirnov test, FDR < 0.05). This demonstrates that even though most of these proteins are not temporally regulated at the gene expression level, their overall gene expression level is still important for cellular proliferation.

### Functional roles in proliferation

If the novel CCD proteins are also involved in the regulation of the cell cycle, loss of function studies should result in cell growth aberrations. To investigate this, we used the Achilles gene essentiality database^50^. This analysis classified 22% (121/541) of the CCD proteins as ‘pan dependent’ genes (i.e., essential for cell survival in >90% of the cell lines screened; Supplementary Table 9). This is a significantly enriched fraction relative to the whole genome (*p* = 5.99×10^−13^, one-sided binomial test) and supports the hypothesis of a functional role of these genes in cellular proliferation. This was true for both novel and known CCD proteins (*p* = 0.0076 and 8.34×10^−18^, one-sided binomial test).

To confirm the role in proliferation, we performed siRNA-mediated gene silencing for a few select novel CCD proteins. Silencing of DUSP18, KLHL38, CD2BP2, and SOX12 decreased cell proliferation rate to 0%, 4%, 12%, and 22% relative to a control, whereas silencing of JPH3 increased cellular proliferation to 170% (Fig. 5D).

Cellular proliferation plays an important role in tumorigenesis, and so we investigated whether expression of the genes encoding CCD proteins are associated with cancer patient outcomes using the Pathology Atlas of HPA^51^, which is based on data from TCGA^52^. Globally, over half of all human genes (54%, 10,577/19,613) were shown to have a prognostic association, where high gene expression were significantly associated with longer survival (favorable) or shorter survival (unfavorable), as previously described^51,52^. Prognostic associations were significantly overrepresented among the cell cycle regulated proteins identified in our study (69%, 376/541 prognostic), supporting the hypothesis of a functional role of these genes in cellular proliferation.

For example, the protein FAM50B is expressed in the nucleus in interphase and translocates to the cytokinetic bridge in mitosis. FAM50B shows a physical interaction with HDAC2, which is also found in the nucleus, where it is involved in the regulation of cell cycle progression^53^. The gene expression survival analysis shows that high FAM50B expression is associated with favorable outcomes in renal cancer, and immunohistochemical (IHC) staining confirms high FAM50B protein expression in normal kidney but low expression in renal cancer (Fig. 5E), suggesting that FAM50B may have an anti-proliferative function.

The protein CD2BP2 was shown to peak in late G2 (Fig. 5E), and its silencing caused decreased cellular growth (Fig. 5D). Higher expression of CD2BP2 had an association with unfavourable prognosis and overall survival in liver cancer. Our IHC analysis confirmed low protein expression in normal liver and high expression in liver tumors (Fig. 5E), suggesting that CD2BP2 expression promotes cell proliferation. We conclude that these novel CCD proteins likely play a functional role in cell proliferation and may have potential to be novel diagnostic or therapeutic cancer targets.

## CONCLUSION

This comprehensive dissection of human cell cycle proteogenomics at the single-cell and subcellular levels is now integrated into the HPA database. We believe it will serve as a valuable resource to accelerate studies towards a greater functional understanding of the human cell cycle, the role of these proteins in tumorigenesis, and identification of novel clinical markers for cellular proliferation.

## Online Methods

### Initial identification of proteins with cell-to-cell heterogeneity

Protein cell-to-cell heterogeneity was identified in the images from the Cell Atlas of the Human Protein Atlas^1^ (HPA v19). Cell-to-cell heterogeneity was identified either as variation in abundance, defined as the change of protein expression levels between single cells within the same field of view, or variations in spatial distribution, defined as translocation of the protein between different subcellular compartments or independent regulation of the protein in two different compartments.

### Cultivation of FUCCI cells

U-2 OS FUCCI cells were kindly provided by Dr. Miyawaki^2^. These cells are endogenously tagged with two fluorescent proteins fused to cell cycle regulators to allow cell cycle monitoring; CDT1 (mKO2-hCdt1+) accumulates in G1 phase, while Geminin (mAG-hGem+) accumulates in S and G2 phases. Cells expressing FUCCI probes are divided into red mKO2(+)mAG(−), yellow mKO2(+)mAG(+), and green mKO2(−)mAG(+) emitting populations. The cells were cultivated in Petri dishes at 37 °C in a 5.0 % CO2 humidified environment in McCoy’s 5A (modified) medium GlutaMAX supplement, (ThermoFisher, 36600021, MA, USA) supplemented with 10% fetal bovine serum (FBS, VWR, Radnor, PA, USA) without antibiotics. The cells were maintained sub-confluent and harvested by trypsinization at log-phase growth (60% confluency) for subsequent analysis.

### Live cell imaging of FUCCI cells

U-2 OS FUCCI cells were grown on a 96-well glass bottom plates (Whatman, Cat# 7716-2370, GE Healthcare, UK, and Greiner Sensoplate Plus, Cat# 655892, Greiner Bio-One, Germany). Approximately 6,000 cells were seeded in the wells and subjected to long-term time-lapse imaging using the molecular device instrument ImageXpress Micro XL (Molecular Device) high content screening equipped with a 20 x Plan Fluor objective and supported with the MetaXpress software, in a 5.0 % CO2 humidified environment at 37 °C. Three Wavelengths were acquired; W1 transmitted light, W2 FITC-3540C filter, W3 CY3-4040C filter. Images were collected every 30 minutes over a course of 72h. For measurement of cell cycle phase length, 8 cells were followed for a cell division cycle, and the time of each phase was calculated as the average for these cells: G1 10.8h; S-transition 2.6h; G2 11.9h with a standard error of the mean of 1.8, 0.125 and 0.225 respectively.

### Antibodies and validation in the Human Protein Atlas

The antibodies used in this study (Supplementary Table 2) were generated within the HPA project. All HPA antibodies are rabbit polyclonal antibodies, designed to bind specifically to as many isoforms of the target protein as possible. Recombinant protein epitope signature tags (PrEST) are used as antigens for immunization^3^. The resulting antibodies were affinity purified using the antigen as affinity ligand^3^. Commercial antibodies were provided by the suppliers and used according to the supplier’s recommendations.

The antibodies are initially quality controlled for sensitivity and lack of cross-reactivity to other proteins, using Western Blot and Protein Arrays^3^. Antibodies that pass the initial quality assessment, with no literature contradicting the results obtained in IHC or IF, are labeled ‘approved’. If there is independent data in UniProt supporting the results in IHC and IF, the antibody is labeled ‘validated’. For enhanced application-specific validation, we use the strategies outlined by the International Working Group for Antibody Validation^4^, including i) validation by gene silencing/knockout, ii) validation by co-staining of FP-tagged protein expressed at endogenous levels, iii) validation by independent antibody targeting a different epitope, iv) capture MS validation^4^. If the antibody has been validated according to the IWGAV guidelines they are labeled ‘enhanced’.

In this study, we included all antibodies that both showed cell-to-cell variability in any cell line in HPA and provided positive IF staining in the U-2 OS cell line in HPA (n = 1,219). We performed extended validation of 157 antibodies with independent antibodies, 29 with gene silencing and 128 with GFP co-staining (Supplementary Table 3). More experimental details on these methods are provided below.

### Protein localization assignment

The distinct localization patterns for each protein in this study are assigned based on prior annotations of the target protein in the HPA Cell Atlas (using the same antibodies) with higher resolution (63x oil confocal imaging)^1^. The staining of all antibodies included in this study were confirmed to match those seen in HPA (automatically in the metacompartment nucleus and/or cytosol; and by manual visual inspection), and thus the localizations were confirmed in this replicated dataset. The enrichment of CCD proteins in subcellular localizations was tested with the binomial one-sided test; the assumptions of this test were satisfied as follows the data is large and nominal, the sample size is less than the population size, the samples are independent, the probability of a given outcome does not not affect the probability of the other.

### Immunocytochemistry

Immunostaining of the U-2 OS FUCCI cells^5^ was performed in 96-well glass bottom plates (Whatman, GE Healthcare, UK, and Greiner Sensoplate Plus, Greiner Bio-One, Germany) coated with 50 μl of 12.5 μg/ml human fibronectin (Sigma Aldrich, Darmstadt, Germany). Approximately 8,000 cells were seeded in each well and incubated at 37 °C for 24 hours. After washing with Phosphate Buffered Saline (PBS, PH=7), cells were fixed with 40 μl 4% ice cold PFA (Sigma Aldrich, Darmstadt, Germany) dissolved in growth medium supplemented with 10% serum for 15 minutes and permeabilized with 40 μl 0.1% Triton X-100 (Sigma Aldrich) in PBS for 3×5 minutes. Rabbit polyclonal HPA antibodies targeting the proteins of interest were dissolved to 2-4 μg/ml in blocking buffer (PBS + 4% FBS) containing 1 μg/ml mouse anti-tubulin (Abcam, ab7291, Cambridge, UK). After washing with PBS, the diluted primary antibodies were added (40 μl/well) and the plates were incubated overnight at 4 °C. After overnight incubation, wells were washed with PBS for 3×10 minutes. Secondary antibodies, goat anti-mouse Alexa405 (A31553, ThermoFisher) and goat anti-rabbit Alexa647 (A21245, ThermoFisher) diluted to 2,5 μg /ml in blocking buffer were added and the plates were incubated for 90 minutes at room temperature. After washing with PBS, all wells were mounted with PBS containing 78 % glycerol before being sealed.

For the incorporation of Bromodeoxyuridine (BrdU) to validate the FUCCI cells, BrdU (10 mg/ml) was diluted to 0.1 mg/ml in fresh pre-warmed growth media. The cells were treated at 37°C for 30 minutes with 40 μl BrdU. Cells were fixed immediately with 40 μl 4% ice cold PFA (Sigma Aldrich, Darmstadt, Germany) dissolved in growth medium supplemented with 10 % serum for 15 minutes and permeabilized with 40 μl 0.1% Triton X-100 (Sigma Aldrich) in PBS for 3×5 minutes. Cells were treated with 1.5 M HCl for 30 minutes. After washing with PBS 3x, the cells were incubated with blocking buffer (PBS + 5% FBS + 0.3% Triton) for 60 minutes. Mouse antibodies targeting the BrdU were dissolved to 2 μg/ml in blocking buffer (PBS + 4% FBS). The diluted primary antibodies were added (40 μl/well) and the plates were incubated overnight at 4 °C. After overnight incubation, wells were washed with PBS for 3×10 minutes. Secondary antibodies, goat anti-mouse Alexa 405 (A31553, ThermoFisher) diluted to 2,5 μg /ml in blocking buffer were added and the plates were incubated for 90 minutes at room temperature. After washing with PBS, all wells were mounted with PBS containing 78 % glycerol before being sealed before imaged using the settings described below. Incorporation of BrdU was confirmed in a subset of the green fucci cells (Extended Data Fig. 3C).

### Image acquisition of FUCCI cell samples

Image acquisition of the immunostained U-2 OS FUCCI cells was performed using ImageXpress Micro XL (Molecular Device) high content screening equipped with a 40 x Plan Apo objective and supported with the MetaXpress software for automated acquisition. Images of the four channels were acquired at room temperature from six positions per sample. Four wavelengths were acquired; W1 for the microtubules DAPI-5060C filter, W2 FITC-3540C filter, W3 CY3-4040C filter and W4 CY5-4040C for the protein of interest. The images were unbinned with a pixel size of 0.1625 × 0.1625 μm. For a limited subset of samples, image acquisition of the immunostained U-2 OS FUCCI cells was performed using InCell 2200 (GE Healthcare) high content screening equipped with a 40 x Plan Apo objective. Images of the four channels were acquired at room temperature from six positions per sample. Four wavelengths were acquired; W1 for the microtubules DAPI (Excitation 390nm/Emission 435nm), W2 FITC (475 nm/511 nm) filter, W3 CY3 (542 nm/597 nm) filter and W4 CY5 (632 nm/679 nm) for the protein of interest. The images were unbinned with a pixel size of 0.1625 × 0.1625 μm.

### Image processing and analysis of images of immunostained FUCCI cells

The segmentation of each cell was performed using the Cell Profiler software^6^, where the overlay of the FUCCI tags was used for nuclei segmentation, and the microtubule staining was used for segmentation of the cell. The mean intensity of the target protein was measured in one of the three main compartments; nucleus, cytosol or cell, based on the a priori-known subcellular localization of the target protein from the HPA Cell Atlas. For each sample, we removed outliers cells with target protein intensities greater than five standard deviations above or below the mean. All image montages were created using Image J and FIJI^6^.

The dataset analyzed in this paper consists of 273,709 cells that exhibited single cell variations during interphase. These were the remaining cells after filtering out-of-focus images, samples with negative antibody staining, samples containing antibodies that were failed in the HPA validation pipeline, samples with very few cells (< 60) due to the difficulty of analyzing cell cycle dependence, cells undergoing mitosis by excluding small nuclei (> 2 standard deviations below mean of nuclei sizes), and segmentation artifacts identified by excluding large nuclei (> 2 standard deviations above mean of nuclei sizes).

To adjust for batch effects observed between plates, which were acquired with two microscopes with different dynamic ranges, the fluorescently tagged CDT1 and GMNN intensities were zero-centered and rescaled in log10 space as follows. (We note that these corrections were not performed for the protein of interest because those measurements are later evaluated on a per-protein basis and not compared between samples like the FUCCI intensities.) The median log10 intensity for CDT1 per plate was subtracted from each log10 intensity on that plate, and the same was done for GMNN intensities. Then, to rescale and normalize the values, the overall minimum log10 intensity for CDT1 was subtracted from all CDT1 intensities, and then those values were divided by the maximum log10 intensity for CDT1; the same was performed for GMNN. The resulting normalized intensity values ranged from 0 to 1 and had batch effects removed due to the centering of each plate around zero for both CDT1 and GMNN in log10 space.

### Fluorescence-Aided Cell Sorting (FACS) of FUCCI cells for single cell RNA-Seq

A total of 1,152 cells were sorted by FACS into three 384-well plates using a BD Influx System (BD Biosciences, San Jose, CA) to allow single-cell RNA-sequencing. We targeted singlet cells using the forward scattering width, and selected approximately equal numbers of cells in G1, S, and G2 phases using the fluorescence intensity of GFP-tagged GMNN (530 nm) and RFP-tagged CDT1 (585 nm). The fluorescence intensities of GMNN and CDT1 were recorded for each cell selected for RNA-Seq that allowed their position within the cell cycle to be determined by pseudotime analysis.

### Single cell RNA-seq analysis

RNA-Seq was collected for each cell using the SMART-seq2 extraction and library preparation protocol^7^ on an Illumina HiSeq 2500 sequencer. FASTQ files (available at GEO SRA under project GSE146773) containing 725 million reads (62.9 +/− 1.7 million reads per cell) of length 43 base pairs were aligned to the human reference genome GRCh38 using the human gene model from Ensembl version 96. The 92 ERCC spike-in sequences^8^ listed in Supplementary Table 11 were appended to the reference genome and gene model before alignment. The alignment and quantification of these data were performed with *RSEM*^9^ (version 1.3.2-0) for quantification using *STAR*^10^ (version 2.7.0f-0) as an aligner. The results used in this analysis were output as TPM values per gene, which combined into a single data frame containing all protein coding genes for all cells using a custom script, and then prepared for downstream analysis using *scanpy*^11^ (version 1.4.4). Cells with fewer than 500 genes detected were filtered (14 cells, 1%) from the analysis, leaving 1,138 cells; genes detected in fewer than 100 cells were filtered (6,547 genes, 33%), leaving 13,450 genes; and TPM values were log transformed. This data was projected onto a *UMAP*^12^ (version 0.3.10), which displayed a cyclical arrangement of cells with five clusters when analyzed by *louvain*^13^ analysis (version 0.6.1). One of these clusters juts out from the circle and is an obvious exception (cluster 5 in Extended Data Fig. 6A). This cluster is enriched for GO terms (analyzed using the tool *genewalk*^14^, version 1.1.0) related to G1 checkpoint and response to DNA damage, and so while these cells are still alive, we believe their proliferation has been arrested due to DNA damage (Extended Data Fig. 6B). This conclusion was corroborated by RNA velocity calculations using *velocyto*^15^ (version 0.17.17) and *scvelo*^16^ (version 0.1.24), which show that these cells are sequestered from cycling (Extended Data Fig. 6A). Therefore, we decided to remove this cluster of cells (117 cells, 10%) from the analysis of cell cycle progression. To evaluate the true positive rate of this analysis, we used a curated list of 97 cell cycle regulating genes from synchronized HeLa cell analysis and multiple single-cell analyses located in Table S5 of a report by Tirosh and colleagues^17^.

### Mock-up bulk analysis of protein cell cycle dependence

For each cell, the GMNN (FUCCI-green) and CDT1 (FUCCI-red) mean intensity values were used to cluster cells into one of three cell cycle clusters using Gaussian clustering. These clusters are denoted as follows: G1: Red(+)Green(−); S-transition: Red(+)Green(+); and G2: (Red(−)Green(+). Please note that the G2 group will also contain some cells in later parts of S-phase. Then, we evaluated whether the median values for the protein intensities were significantly different between the three groups using the two-sided Kruskal-Wallis statistical test^16^ (the samples were random, mutually independent, continuous, and the distributions were similar for intensities within cell cycle phases measured with the same antibody). Resulting FDR values less than 0.01 were considered significant in this mock-up bulk analysis (Supplementary Table 2). Boxplots were used to illustrate these results within Fig. 1 that were generated with the default elements in MatPlotLib^19^ (center line, median; box limits, first quartile (Q1) and third quartile (Q3); whiskers, Q1 − 1.5 × IQR and Q3 + 1.5 × IQR, where IQR is the interquartile range; points, outliers).

### Polar-coordinate pseudotime model

In this work, we utilized the FUCCI system to model cell cycle position. To generate a continuous representation of cell cycle position, we utilized a polar regression based on a log-scale scatter plot of GMNN (FUCCI-green) and CDT1 (FUCCI-red), where each point represents a single cell (Extended Data Fig. 3). This data was shifted, such that the origin point lay at the center of mass. This allowed us to use the fractional radius of the circle to estimate time for each cell as traced by a ray from the origin generating a polar regression representing continuous cell cycle position. The cell-division point was selected by using the area of lowest cell density on the polar ray from the origin. This is justified by the knowledge that the M phase, where cells express neither GMNN nor CDT1 highly, is much shorter than all other phases. The selected point was validated via visual inspection of nearby cells. This allowed us to linearize the progression of time from 0 to 1 representing the fractional distance along this polar axis from 0 to 360 degrees.

### Moving average model

Cell-cycle correlation was measured using a moving average model within the linearized time from the polar fit described above. A range of window sizes were tested from 5-30 cells for protein analysis and 50-150 cells for RNA analysis. The analysis proved robust to these ranges of window sizes, and results reported are for window sizes of 10 cells for protein analysis and 100 cells for RNA analysis, which were chosen to balance the robustness to outliers with potentially destroying signal. We plotted the results of the moving average models for various genes in Figs. 1, 3, 4, and 5 with protein in blue and RNA in orange: line representing the moving mean, darker color representing 25th to 75th percentile range, lighter color representing 10th to 90th percentile range, and points representing the individual cell data.

### Percent explained variance

We used the metric of percent explained variance to describe the quality of our model fit. This metric is appealing because it is scale-invariant. That is, unlike a *p*-value significance metric, which becomes more significant as sample size increases, the percent-variance converges to a stable solution as more cells are sampled. The percent explained variance is calculated as:

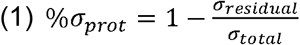

Here, σ_*prot*_ represents the variance of the protein of interest for an experiment and σ_*residual*_ represents the variance remaining calculated from the moving average line along the pseudo-time axis.

A high percent variance value is observed for both a sample with cell cycle dependence and for a sample with a highly fluctuating moving average, and so we tested whether this value was reflective of cell cycle dependence using a randomization analysis. The percent variance was calculated using the original ordering of cells, as well as for 10,000 permutations of the cells in the protein and RNA analyses. We used one-sided Wilcoxon tests of whether the percent variance explained by the cell cycle for each gene was significantly higher (*p* < 0.01; the data met the assumptions of the Wilcoxon test: the permutations were independent and the difference between permuted percent variances and the original percent variance is continuous and ordinal) than the permuted orderings for protein and RNA, corrected for multiple testing using the Bonferroni correction; we also applied an additional cutoff to the additional percent variance explained than random (Extended Data Fig. 6D). Specifically, an additional 8% percent variance than random was required for a protein or RNA to be called significantly cell cycle dependent (CCD). Examples of permutations leading to the minimum, first quartile, median, third quartile and max percent variance are provided in Extended Data Fig. 6E. These results can be viewed for all genes in the scatter plots of percent variance and Gini index in Fig. 2A for protein and Extended Data Fig. 6C for RNA. We report a small overlap of CCD proteins and CCD transcripts (15%); while single-cell sequencing has a high dropout rate, which may contribute false negative CCD transcripts, this only accounts for only 12% of the negatives in this dataset (filtered or dropout transcripts were observed for 146/1,253 proteins analyzed in interphase and 38/326 proteins found to be CCD in interphase), and thus dropouts cannot account for the small overlap observed.

### Distributions of protein expression among cell populations

The patterns of variability across the measured population of cells were investigated for each of the 1,219 proteins. k-means clustering was performed using the kurtosis and skewness features of the distribution of the mean intensity per cell for each protein. Each gene was assigned to a specific cluster, and the optimal number of clusters was chosen using the Elbow method. Three distinct clusters were found. Cluster 1, the largest cluster (n=1,030), contained 90% (291/322) of CCD proteins and 85% (739/868) of non-CCD proteins. The lower segment of Cluster 1 contained proteins such as GATA6 and CCNB1 (Extended Data Fig. 2C). Cluster 2, the second largest cluster (n=148), contained proteins with slightly skewed distribution profiles with a sharp peak distribution, as exemplified by DEF6 (Extended Data Fig. 2C). This cluster contained 10% (31/322) of the ccd proteins and 14% (118/868) of non CCD proteins. Cluster 3 (n=12) contained only non-CCD proteins, where the variation was highly skewed and tailed with few cells expressing the protein. We also determined that 992 proteins (81%) had unimodal and 227 proteins (19%) had bimodal population intensity distributions (Extended Data Fig. 2D) as exemplified by SLC25A42 (Extended Data Fig. 2F) and HPSE (Extended Data Fig. 2G). These results show that cell cycle dependent variations are mostly unimodal with a normal distribution across a log-phase growing population of cells.

Gaussian clustering was used to allow statistical determination if populations of cells were unimodal or bimodal for the expression of each protein. Specifically, we aimed to analyze cell cycle dependence in samples that had two distinct populations of cells, high-and low-expressing for the protein of interest. If the population was bimodal, we asked whether the protein was CCD in either population. We determined that 992 proteins (81%) had unimodal intensity distributions, and 227 proteins (19%) were found to display bimodality by requiring a significant difference in intensity between the two clusters (FDR < 0.01 by two-sided Kruskal-Wallis test^18^, adjusted by Benjimini-Hochberg correction^20^; the samples were random, mutually independent, continuous, and the distributions were similar for intensities within each cluster measured with the same antibody) and a fold change of 2 (Extended Data Fig. 2D). Some CCD proteins display strong bimodality, such as GATA6 (Extended Data Fig. 2E), and so before evaluating the cell cycle dependence of the two populations, we required there to be no significant difference in pseudotime between the two clusters (FDR > 0.01 by two-sided Kruskal-Wallis test^18^, adjusted by Benjimini-Hochberg correction^20^; the samples were random, mutually independent, continuous, and the distributions were similar for pseudotimes within each cluster measured with the same antibody). We required there to be at least 50 cells in each cluster to allow for CCD analysis, and statistical tests assessing CCD for distinct high- and low-expressing cell populations were concatenated onto tests of unimodal populations for multiple testing corrections. Of the 43 bimodal samples that were analyzed for cell cycle dependence, 12 were CCD in one cluster (2 of these were CCD when analyzed unimodally), and 2 were CCD in both clusters (both were also CCD when analyzed unimodally). The protein SLC25A2 provides a CCD example in one cluster at lower intensities, where we observe second harmonic cycling (Extended Data Fig. 2F). We observed non-CCD patterns for both high and low expressing cells for the remaining 29 proteins displaying bimodal protein expression, including HPSE (Extended Data Fig. 2G).

### Gene set enrichment analysis

Functional enrichment analysis for the GO domain biological process was performed using the Database for Annotation, Visualization and Integrative Discovery (DAVID) tool^21^ and Cytoscape^22^ v3.6.1 was used for the network visualization. The enrichment map Cytoscape plugin was used to visualize the results of the highly significant gene-set enrichment as a network^23^. Functional enrichment of variably expressed metabolic enzymes was performed using GOrilla^24^ using cutoffs of *p* < 0.001, n > 4 genes, and 2X enrichment. A list of proteins with a known function related to the cell cycle was obtained by combining all hits related to the cell cycle or cell proliferation from the databases Cyclebase^25^ v3.0; Reactome^26^ and QuickGO^27^.

### Protein interaction analysis

The interaction analysis was done using the Search Tool for the Retrieval of Interacting Genes/Proteins (STRING) database^28^ v10.5, where a medium confidence (0.4) score was used to highlight the protein-protein interaction edges.

### Bulk RNA extraction and RNA sequencing

The RNA extraction and sequencing were performed as previously reported^1,29,30^. Tissue samples were embedded in Optimal Cutting Temperature (OCT, Sakura) compound and stored at –80°C. HE-stained frozen sections (4 μm) were prepared from each sample using a cryostat and the CryoJane® Tape-Transfer System (Instrumedics, St. Louis, MO, USA). Three sections (10 μm) were cut from each frozen tissue block and collected in a tube for subsequent RNA extraction 108. Total RNA was extracted from the cell lines and tissue samples using the RNeasy Mini Kit (Qiagen, Hilden, Germany) according to the manufacturer’s instructions. Only samples of high-quality RNA (RNA Integrity Number ≥ 7.5) were used in the following mRNA sample preparation for sequencing.

A total of 172 samples from 37 tissues and organs was sequenced using Illumina HiSeq 2000 and HiSeq 2500, and the standard Illumina RNAseq protocol with a read length of 2×100 bases. Briefly, the reads were mapped to the human genome (GRCh37) using Tophat^31^ v2.0.8b. Transcript abundance estimation was performed using Kallisto^32^ v0.42.4. For each gene, the abundance was reported in ‘Transcript Per Million’ (TPM) as the sum of the TPM values of all its protein-coding transcripts. For each cell line and tissue type, the average TPM value for replicate samples was used as an abundance score.

### Co-Expression network analysis

The co-expression networks for different tissues and cancers were downloaded from TCSBN database^33^. The nodes (genes) in the networks were classified into three categories: i) candidate CCD genes (T1), ii) known CCD genes (T2) and iii) other genes (T3). Following that, the shortest path in the co-expression network was compared between each category by using a simple Breadth-First Search (BFS) method. The distribution between shortest path of T1-T2 was compared with T3-T2 by FDR-Adjusted Kolmogorov-Smirnov one-sided test (FDR < 0.05; the samples satisfied the assumptions of the test: samples of lengths were random, and the theoretical distribution of lengths is continuous and fully defined). The aforementioned analyses were performed using Python, with Scipy module^34^ for the statistical analysis and Igraph^35^ for the network analysis and manipulation.

### Analyses of PTM occupancy with MetaMorpheus

We mined a deep bottom-up proteomics dataset collected using U-2 OS cells for abundant post-translational modifications (PTMs) in bulk^36^ and for phosphorylations in phospho-enriched U-2 OS cells^37^ using the Global PTM Discovery (G-PTM-D) search strategy^38^ with MetaMorpheus^39^ (version 302; https://github.com/smith-chem-wisc/MetaMorpheus). This strategy uses a two stage search that first looks for candidate PTMs using characteristic mass shifts, and then identifies peptides with possibly new PTM sites using a narrow mass search that allows strict controls of false discoveries.

The bulk proteomic analysis (bulk) and phosphoproteomic analysis (phospho) were performed separately and yielded complementary information. The first search added 38,877 (bulk) and 215,535 (phospho) candidate PTM sites to the protein database by finding peptide spectral matches at mass differences characteristic of PTMs. For the phosphoproteomic analysis, 54% of these candidate PTM sites were phosphorylations compared to 4% of candidate PTM sites in bulk analysis. The subsequent narrow mass tolerance search identified 395,583 PSMs (bulk) and 135,043 PSMs (phospho) within 1% FDR, 66,684 peptide base sequences (bulk) and 36,366 peptide base sequences (phospho) within 1% FDR, and 76,324 peptides (bulk) and 49,164 peptides (phospho) including modified forms within 1% FDR. PTM site occupancies were calculated by dividing the counts of modified peptides by the total peptides crossing the site, as recorded in the ProteinGroups.tsv file output by MetaMorpheus. Of the 9,848 proteins (bulk) and 11,228 proteins (phospho) detected, 2,573 proteins (bulk, 26%) and 11,057 proteins (phospho, 98%) were detected with modifications. The modified proteins studied in this analysis had higher coverage (34%, bulk; 31%, phospho) than proteins on average (18%, bulk; 30% phospho). Metal adducts and sample preparation artifacts were included in G-PTM-D and the final search, but were excluded from the PTM occupancy analysis. Boxplots for Fig. 3E-F illustrating PTM occupancy and protein melting point distributions were constructed as follows: center line, median; box limits, Q1 and Q3; whiskers, 1.5X IQR; n.s., not significant, *p<0.05; **p<0.01; ***p<0.001.

We note that bulk proteomic and phosphoproteomic methodologies have different limitations. Bulk PTM analysis may miss low stoichiometry PTMs and those specific to rarer cell types, like mitotic cells. For example, tightly controlled PTMs specific to mitosis may be missed, since they would likely fall below the detection limit of MS with only ~2% of cells in log phase growth undergoing mitosis. While phosphoproteomic analysis may enable detection of low stoichiometry phosphorylations and those that occur in rarer cell types, phosphoproteomic analysis may miss highly abundant PTMs that are seen in bulk PTM analysis.

The following settings were used for both the G-PTM-D and narrow mass searches: protease = trypsin; maximum missed cleavages = 2; minimum peptide length = 7; maximum peptide length = unspecified; initiator methionine behavior = Variable; max modification isoforms = 1024; fixed modifications = Carbamidomethyl on C and on U; variable modifications = Oxidation on M; product mass tolerance = ± 20 PPM. For the phosphoproteomic analysis, additional fixed modifications, TMT6-plex on X and TMT6-plex on K were specified, since the dataset used 6-plex TMT labeling, and the data were searched with common contaminant proteins that were removed from the final results. Data files were processed on a Dell Inc. Precision Tower 7910 computer using Windows version 6.2.9200.0 with a 64-Bit processor and 24 cores operating at 2.993GHz and 128GB installed RAM. The narrow mass search was performed with the custom database output by G-PTM-D using a simple precursor mass tolerance of ± 5 PPM around 0 Da. The G-PTM-D search used a UniProt-XML database downloaded on Apr 18, 2019 and used precursor mass tolerances of ± 5 PPM around −74.048012822, −28.031300129, −18.010564684, −17.026549101, −16.042533, −14.015650065, −2.01565, −1.031634, 0, 0.984015583, 1.968032, 14.015650064, 14.999666, 15.99491462, 16.978931, 27.010899036, 27.99491462, 28.031300129, 28.990163592, 31.98983, 42.010564684, 42.046950193, 42.994581, 43.005813656, 43.989829239, 44.985078211, 68.026214748, 70.041864813, 79.956815033, 79.966330889, 86.000393923, 87.044604465, 100.016043988, 114.031694052, 159.932662, 203.079372521, 229.014009359, and 541.06110975 Da.

### Immunohistochemical staining

Immunohistochemical (IHC) staining of tissue microarray (TMA) sections and slide scanning were performed as previously described^40^. Note that both the normal and cancer tissues were stained simultaneously with identical conditions and imaged with identical settings to ensure that the obtained staining intensities are directly comparable.

### Validation of antibodies and functional role in proliferation by gene silencing

U-2 OS FUCCI cells were cultivated as described above. The cells were seeded in 96-well glass bottom plates (Whatman, GE Healthcare, UK, and Greiner Sensoplate Plus, Greiner Bio-One, Germany) coated with 50 μl of 12.5 μg/ml human fibronectin (Sigma Aldrich, Darmstadt, Germany). Approximately 6,000 cells were seeded in each well and incubated at 37 °C for 24 hours. Small interfering RNA (siRNA) oligonucleotides were obtained from Life Technologies (Silencer Select 0.1nm, Supplementary Table 3). Cells were transfected 24 h after seeding (at 60% confluency) with 10nM oligonucleotide using lipofectamine. After 24h, the media with the transfection reagents were replaced with fresh media. Thereafter, the cells were fixed, stained and imaged with InCell 2200 as described above. Image processing and analysis of images of immunostained FUCCI cells was applied also as described above, where the mean intensity signal of the target protein was compared between control and siRNA. Boxplots were generated with the default elements in R studio v1.1.423 (center line, median; box limits, Q1 and Q3; whiskers, Q1 − 1.5 × IQR and Q3 + 1.5 × IQR; points, outliers). To assess the cellular proliferation rate we compared the proliferation to the control cells.

### mNG-tagged cell line generation and cultivation

We applied a high-throughput CRISPR/Cas9 mediated approach^41,42^ for endogenous mNG gene tagging at the N- or C-terminus. Cas9/sgRNA ribonucleo-protein complexes were prepared by assembling 100 pmol Cas9 protein and 130 pmol sgRNA. sgRNA (Supplementary Table 12) was diluted in 6.5 μl Cas9 buffer (150 mM KCl, 20 mM Tris pH 7.5, 1 mM TCEP-Cl, 1 mM MgCL_2_, 10% glycerol) and incubated at 70 °C for 5 min. After addition of the Cas9 protein, the RNP assembly was performed at 37 °C for 10 min. 1 μl of HDR template was added to a total volume of 10 μl. Hek293T^mNG1-10^ cells were treated with 200 ng/mL nocadozole (Sigma) for 16 hours prior to electroporation which was carried out according to the manufacturer’s instructions using SF-cell line reagents (Lonza) in an Amaxa 96-well shuttle Nucleofector device (Lonza). Cells were washed with PBS and resuspended to 10^4^ cells/μl in SF solution. Around 2x×10^5^ cells were mixed with the Cas9/sgRNA RNP complexes, electroporated using the CM-130 program and rescued in cell culture medium in a 96-well plate. Electroporated cells were cultured and maintained for 8 days prior to FACS analysis. All synthetic nucleic acid reagents (Table S4) were ordered from Integrative DNA Technologies (IDT DNA).

Endogenously tagged HEK293T cells were cultured in high glucose Dulbecco’s Modified Eagle Medium, supplemented with 10 % FBS, 1 mM glutamine and 100 μg/ml penicillin/streptomycin (Gibco). Cells were maintained in 96-well plates at 37 °C in a 5.0 % CO2 humidified environment.

### Fluorescence activated cell sorting (FACS) and polyclonal enrichment

We performed analytical flow cytometry on a SH800 instrument (Sony) to assess the tagging efficiency of the cell lines. Polyclonal populations of the 0.5% highest mNG-expressing cells were established for follow-up studies.

### Generation of Illumina amplicon sequencing libraries

DNA repair outcomes were characterized by Illumina amplicon sequencing as an important quality control step. Hek293T^mNG1-10^ cells were grown in a 96-well plate at around 80% confluency. Cells were washed with PBS and thoroughly resuspended in 50 μl QuickExtract (Lucigen) and incubated at 65 °C for 20 min and 98 °C for 5 min. 2 μl gDNA, 20 μl 2x KAPA, 1.6 μl of 50 μM forward and reverse primer (Supplementary Table 13), 8 μl 5 M betaine and 8.4 μ H2O were run using the following thermocycler settings: 3 min at 95 °C followed by 3 cycles of 20 s at 98 °C, 15 s at 63 °C 20 s at 72 °C, 3 cycles of 20 s at 98 °C, 15 s at 65 °C 20 s at 72 °C, 3 cycles of 20 s at 98 °C, 15 s at 67 °C 20 s at 72 °C and 17 cycles of 20 s at 98 °C, 15 s at 69°C 20 s at 72 °C. Unique index sequences for multiplexed sequencing were attached to each amplicon in a barcoding PCR. 1 μl of 2 nM PCR product, 4 μl of forward and reverse indexed barcoding primer, 20 μl 2x Kapa and 11 μl H2O were run with the following thermocycler settings: 3 min at 95 °C, 10 cycles of 20 s at 98 °C, 15 s at 68 °C, 12 s at 72 °C. Product concentrations in 200 – 600 bp range were quantified using a Fragment Analyzer (Agilent) and pooled at 500 nM. Amplicon sequencing library was purified using a Dual Solid Phase Reversible Immobilization at a 0.6 and 1.1x bead/sample ratio. Sequencing was performed using an Illumina MiSeq system at the CZ Biohub Sequencing facility. Sequencing outcomes were characterized using CRISPResso^43^.

### Spinning-disk confocal timelapse acquisition of mNG tagged cells

Around 10,000 endogenously tagged HEK293T cells were grown on a fibronectin (Roche)-coated 96-well glass bottom plate (Cellvis) for 24 hours. Cells were counterstained in 0.05 μg/ml Hoechst 33342 (Thermo) for 30 min at 37 °C and imaged in complete DMEM without phenol-red. Live-cell imaging was performed at 37 °C and 5% CO2 on a Dragonfly spinning-disk confocal microscope (Andor) equipped with a 1.45 N/A 63x oil objective and an iXon Ultra 888 EMCCD camera (Andor). Images were acquired every hour for 40-50 hours.

### Antibody validation in mNG tagged cell lines

To validate a subset of the antibodies in this study, 20,000 endogenously tagged HEK293T cells were seeded per well on a fibronectin (Roche)-coated 96-well glass bottom plate (Cellvis) and cultivated for 24 hours. The mNG tagged cells were fixed in 4% formaldehyde (Thermo Scientific, 28908) and immunostained with the corresponding antibody as described before.

## Acknowledgements

We acknowledge the entire staff of the Human Protein Atlas program. We acknowledge Dr. Hisao Masai (Tokyo Metropolitan Institute of Medical Science) for providing the stable U-2 OS FUCCI cell line. We also acknowledge Dr. Magdalena Otrocka from the National Laboratories for Chemical Biology at Karolinska Institutet (LCBKI) for access to imaging infrastructure, Dr. Petter Ranefall and Dr. Carolina Wählby for providing support for establishing the Cell Profiler pipeline, and Dr. Lloyd M. Smith for providing access to computational resources. Funding was provided by the Knut and Alice Wallenberg Foundation (2016.0204) and Swedish Research Council (2017-05327) to E.L.

## Author Contributions

E.L. conceived the study. D.M., D.P.S and E.L. developed the methodology for the study. D.M., L.S., R.S., C.G, and P.T. carried out the immunofluorescent experimental work and contributed to the cell atlas implementation. C.G., N.H.C., and M.D.L, performed the GFP tagging and analysis. D.M, A.B and C.S performed the siRNA antibody validation and growth assays. D.M., A.J.C, D.P.S. and E.L. carried out protein imaging data analysis and investigation. A.J.C. performed single-cell RNA-Seq, MS proteomic, and proteogenomic analyses. F.D. analyzed the bulk RNA-seq data. M.A., C.Z. and A.M. carried out the co-expression network analysis. C.L. and F.P. provided the tissue data. D.M., A.J.C and E.L. wrote the manuscript. B.A., D.P.S, O.C and P.T revised the manuscript. D.M., A.J.C., C.G and D.P.S. created the figures. M.U. initiated the HPA project and provided antibodies. E.L. supervised and administered the project and acquired funding. All authors reviewed and approved the final manuscript before submission.

## Supplementary Information

Supplementary information is available for this paper.

## Corresponding Author

Correspondence and requests for materials should be addressed to E.L.

## Life Sciences Reporting Summary

Further information on experimental design is available in the Nature Research Reporting Summary linked to this article.

## Data Availability Statement

The images from the Human Protein Atlas are available at: https://www.proteinatlas.org. The images from the FUCCI screening are available as jpg in the Human Protein Atlas (v20 released upon final publication). Uncompressed images are available upon request. The RNA-sequencing data is available at www.ebi.ac.uk/arrayexpress/experiments/E-MTAB-2836/. The single-cell RNA-Seq data is available at GSE146773.

The data is available for download in the supplementary material. Images and Cell Atlas transcriptome and proteome data along with interpretation and classification is available in the Human Protein Atlas (www.proteinatlas.org/humancell).

## Code Availability Statement

The cell profiler pipeline for image analysis and the code for generating the polar-coordinate pseudotime model are available at: https://github.com/CellProfiling/fucci_screen

## Extended Data Figure legends

**Extended Data Fig. 1:** Validation examples of CCD and non-CCD proteins using co-localization with mNG tagged proteins and siRNA gene silencing.

A) The specificity of several dozen antibodies (45 in total with 10 examples presented here; Supplementary Table 3) targeting proteins with CCD expression was validated by co-localization with mNG-tagged protein. Scale bars correspond to 10 μm.

B) The specificity of several dozen antibodies (79 in total with 10 examples presented here; Supplementary Table 3) targeting proteins with non-CCD variable expression was validated by co-localization with mNG-tagged protein. Scale bars correspond to 10 μm.

C) The specificities of 29 antibodies (two presented here; Fig. 1C; Fig. 4B; Fig. 5D; Supplementary Table 3) were validated with siRNA-mediated gene silencing that resulted in significantly lower staining intensity. Scale bars correspond to 10 μm.

**Extended Data Fig. 2:** Variation distribution, bimodality, and cell cycle dependence in distinct high-and low-expressing cell populations.

A) Scatterplot showing the three clusters generated by k-mean clustering based on kurtosis and skewness as features for CCD proteins.

B) Scatterplot showing the three clusters generated by k-mean clustering based on kurtosis and skewness as features for non-CCD proteins.

C) Violin-plots and histograms showing the population distributions of the normalized mean intensity of each cell per protein for three selected CCD examples (GATA6; CCNB1 and DEF6).

D) Bimodal protein distributions were evaluated for cell cycle dependence separately in both low- and high-expressing cells if the two populations were determined to be distinct. This determination was performed using a Kruskal-Wallis test, adjusted for multiple testing, and if they had greater than a 2-fold difference in expression between them.

E) GATA6 expression over the cell cycle. While GATA6 produces a bimodal population intensity distribution, it represents a single population of cells and was evaluated for cell cycle dependence as such rather than as two populations of high- and low-expressing cells.

F) SLC25A42 expression over the cell cycle is exhibited as two distinct populations of high- and low-expressing cells (left). The low-expressing cells (center) have CCD expression, forming a second harmonic over the cell cycle, while the high-expressing cells (right) do not display correlation to the cell cycle.

G) HPSE expression over the cell cycle has two distinct populations of high- and low-expressing cells that both display non-CCD expression.

**Extended Data Fig. 3:** Validation of the FUCCI cell model.

A) Images from a time-lapse microscopy series of FUCCI cells show that the length of the cell cycle phases marked by the different colours of the FUCCI tags are 10.8 hrs for G1 10.8h, 2.6 hrs for S-transition, and 11.9 hrs for G2. U-2 OS FUCCI cells allow monitoring the cell cycle by expressing two fluorescently-tagged cell cycle markers: CDT1 during G1 phase (red), GMNN during S and G2 phases (green), and their co-expression during the S-transition (yellow). Scale bars correspond to 10 μm.

B) The polar coordinate model transfers the FUCCI marker information into a linear model of pseudo-time. The tagged CDT1 (red) and GMNN (green) log-intensities are displayed across pseudotime that they are used to calculate. The solid line is the moving average across 100 cells (273,709 total), and light shading fills between the 25th and 75th percentiles of intensity.

C) BrdU incorporation (blue) into the FUCCI cells. Incorporation of the synthetic nucleotide BrdU occurs during the DNA replication in S-phase of proliferating cells and remains through the rest of the cell cycle. Here G1 (red) cells show no incorporation of Brdu, whereas most of the cells with green nuclei show incorporation of Brdu which is indicative of the S and G2 phases. These results show the validity of the FUCCI model.

**Extended Data Fig. 4:** Validation of CCD and non-CCD protein using mNG-tagged live cell imaging.

A) The specificity of the anti-CPSF6 antibody by co-localization with mNG-tagged CPSF6. Scale bars correspond to 10 μm.

B) The specificity of the anti-ANLN antibody by co-localization with mNG-tagged ANLN. Scale bars correspond to 10 μm.

C) Timelapse microscopy for mNG-tagged ANLN protein demonstrates an increase in nuclear expression over the course of the cell cycle with a peak in late G2. During mitosis and cytokinesis, ANLN localizes to the cell membrane and is involved in the formation of the cleavage furrow (t=17-18 hours and t=35 hours). Scale bar corresponds to 10 μm.

**Extended Data Fig. 5:** Validation of variability observed with antibody staining

A) Box plot showing gene expression levels for all proteins that exhibit cell-to-cell heterogeneity and all proteins in the HPA Cell Atlas (mapped proteome). There is no significant difference (*p* > 0.01) of gene expression levels between the antibodies included in this study and all antibodies used in the HPA Cell Atlas (*p* = 0.03, two-sample Student’s *t*-test). Center line, median; box limits, first quartile (Q1) and third quartile (Q3); whiskers, Q1 − 1.5 × IQR and Q3 + 1.5 × IQR, where IQR is the interquartile range; points, outliers.

B) Bar plot showing staining intensity levels for antibodies included in this study and all proteins in the HPA Cell Atlas (mapped proteome). The average immunofluorescence signal intensity measured for proteins showing cell-to-cell variation and for all proteins in the HPA Cell Atlas shows no significant difference for low signal antibodies. Low signal antibodies are not enriched among the cell-to-cell variability dataset (*p* = 0.99, Binomial test).

C) Since the HPA antibodies are purified on affinity columns coupled with their corresponding antigen, antibody concentration after purification can serve as a proxy for antibody affinity (albeit not a perfect proxy). Box plot showing antibody concentration levels for all proteins that showed cell-to-cell heterogeneity and all proteins in the HPA Cell Atlas (mapped proteome; center line, median; box limits, first quartile (Q1) and third quartile (Q3); whiskers, Q1 − 1.5 × IQR and Q3 + 1.5 × IQR; points, outliers). There is no significant difference between the antibodies included in this study and all antibodies used in the HPA Cell Atlas, hence we can conclude that the cell-to-cell heterogeneity is most likely not due to differences in antibody affinity. The average antibody concentration for the antibodies published on the HPA Cell Atlas is 0.1710 mg/ml, and the average concentration for the antibodies used in this study is 0.1712 mg/ml (*p* = 0.1084, two-sample Student’s *t*-test).

D) Box plot showing the variance for all proteins that showed cell-to-cell heterogeneity and the microtubules (center line, median; box limits, first quartile (Q1) and third quartile (Q3); whiskers, Q1 − 1.5 × IQR and Q3 + 1.5 × IQR). Proteins showing heterogeneity show significantly higher variance than the variance of the microtubules measured from each well (*p* = 9.7×10^−304^, one-sided Kruskal-Wallis test).

E) Box plot showing the Gini index for all proteins that showed cell-to-cell heterogeneity and for microtubules (center line, median; box limits, first quartile (Q1) and third quartile (Q3); whiskers, Q1 − 1.5 × IQR and Q3 + 1.5 × IQR). Proteins displaying heterogeneity show significantly higher Gini indexes than microtubules (*p* < 0.0, one-sided Kruskal-Wallis test).

**Extended Data Fig. 6:** Filtering of non-cycling cells for RNA-Seq analysis and illustrations of randomization analysis.

A) Dimensionality reduction of single-cell RNA sequencing data by UMAP with RNA velocities overlayed in arrows and with coloring based on clustering. Each point represents a cell, and cluster 5 (brown) appears to be cells that are sequestered from cycling after G1 (top left).

B) Gene ontology biological process terms for cluster 5 show enrichment for terms that indicate arrest of cell cycling after G1, such as “mitotic G1 DNA damage checkpoint,” and so the cells in cluster 5 were removed from the main analysis.

C) Percent variance explained for RNA is shown against the Gini index for the gene (blue: CCD, red: non-CCD).

D) Randomization analysis of the protein IF data (left) and single-cell RNA-Seq data (right) for each gene was used to determine if a protein or RNA was CCD (blue) or non-CCD (red). The significance scores, adjusted for multiple correction, on the vertical axis show that nearly all proteins and RNAs are significant, and so requiring 8% additional percent variance explained by the cell cycle than random was the predominant cutoff.

E) Examples of NFAT5 protein IF data (blue points and trace) and randomizations of cell order in pseudotime (red points and trace). These examples provided (from left to right) produced the minimum, first quartile, median, third quartile, and maximum percent variance explained by the cell cycle.

**Extended Data Fig. 7:** Time lapses of cell cycle regulated proteins.

A) The specificity of the anti-AURKB antibody was validated by co-localization with mNG-tagged AURKB. Scale bars correspond to 10 μm.

B) Time lapse microscopy for mNG-tagged AURKB protein demonstrates an increase in protein expression over the course of the cell cycle with a peak in late G2. During mitosis, AURKB localizes to the mitotic chromosomes (t=13 hours and t=29 hours). Scale bars correspond to 10 μm.

C) Time lapse microscopy for mNG-tagged KIFC protein demonstrates a CCD translocation from the nucleoplasm to the kinetochores in late G2 into mitosis (t=14 hours and t=30 hours) and early G1. Scale bars correspond to 10 μm.

D) The specificity of the anti-GATAD1 antibody was validated by co-localization with mNG-tagged GATAD1. Scale bars correspond to 10 μm.

E) Time lapse microscopy for mNG-tagged GATAD1 protein demonstrates a decrease in nuclear abundance throughout the cell cycle (mitosis at t=35 hours). Scale bars correspond to 10 μm.

F) Time lapse microscopy for mNG-tagged NET1 protein demonstrates a CCD increase of protein abundance and a translocation from the nucleoplasm to the nucleolus (mitosis at t=35 hours). Scale bars correspond to 10 μm.

**Extended Data Fig. 8:** Clustering of gene expression in tissue and tumors for known and novel CCD proteins.

A) Hierarchical clustering of bulk transcript expression in various normal and cancer tissue types (log-transformed TPM values) for known CCD proteins. The expression levels of the proliferation markers MCM6, CDK1, PCNA, MCM2 and KI67 are highlighted on top as a general measure of the proliferative activity of the tissues. Four clusters are identified: (c1) and (c2) contain mostly normal tissues with midrange expression of the proliferation markers; (c3) contains tissues with high expression of the proliferation markers, including tumors; and (c4) contains normal tissues with low proliferative activity.

B) Box plots displaying the average transcript level for known CCD proteins in the four different clusters from panel A). There is a significant difference of gene expression levels between the different clusters (*p* < 0.01 between all the clusters, one-sided Kruskal-Wallis test; center line, median; box limits, first quartile (Q1) and third quartile (Q3); whiskers, Q1 − 1.5 × IQR and Q3 + 1.5 × IQR; points, outliers).

C) Hierarchical clustering of bulk transcript expression in various normal and cancer tissue types (log-transformed TPM values) for novel CCD proteins. The expression levels of the proliferation markers MCM6, CDK1, PCNA, MCM2 and KI67 are highlighted on top as a general measure of the proliferative activity of the tissues. Four clusters are identified: (c1) contains normal tissues with midrange expression of the proliferation markers and tissues with high expression of the proliferation markers, including tumors; (c2) contains bone marrow and bone marrow cancer tissues; (c3) contains cerebral tissues with testis; and (c4) contains normal tissues with low proliferative activity.

D) Box plots displaying the average transcript level for novel CCD proteins in the four different clusters from panel C). There is a significant difference between gene expression in the different clusters except between clusters c2 and c4 (*p* < 0.01 between all the clusters except for cluster c2 and c4, where *p* = 0.3, one sided Kruskal-Wallis test; center line, median; box limits, first quartile (Q1) and third quartile (Q3); whiskers, Q1 −1.5 × IQR and Q3 + 1.5 × IQR; points, outliers).

## Supplementary information table titles

Supplementary Table 1: List of all proteins in HPA that show cell-to-cell variability. Supplementary Table 2: Protein temporal data

Supplementary Table 3: Antibody validation: mNG, siRNA, different epitopes Supplementary Table 4: Proteins in mitotic structures

Supplementary Table 5: RNA temporal data

Supplementary Table 6: PTM site occupancies from bulk MS proteomic study of U2OS. Supplementary Table 7: PTM site occupancies from phospho-enrichment MS proteomic study of U2OS.

Supplementary Table 8: Average melting point data for proteins Supplementary Table 9: Gene essentiality analysis

Supplementary Table 10: ERCC spike in sequences for single-cell RNA-seq analysis. Supplementary Table 11: CRISPR gRNA sequences.

Supplementary Table 12: Sequence of primers used for amplicon library preparation.

